# Shading Reduces Water Deficit in Strawberry (*Fragaria ×ananassa* Duch.) Plants during Vegetative Growth

**DOI:** 10.1101/2021.08.05.455240

**Authors:** Henry Alexander Cordoba-Novoa, Maria Mercedes Pérez, Brahyam Emmanuel Cruz, Nixon Flórez, Stanislav Magnitskiy, Liz Patricia Moreno

## Abstract

Strawberry (*Fragaria ×ananassa* Duch.) is a commercially important crop with high water requirements, for which it is necessary to find strategies that mitigate the influence of water deficit on plant growth. This study was aimed to evaluate the effects of shading on the vegetative growth of strawberry cv. Sweet Ann under water deficit. The treatments consisted of the combination of two levels of shading (light intensity reduced on 47% vs. non-shaded plants) and two levels of water availability (water deficit vs. well-watered plants). The water deficit reduced the leaf water potential from −1.52 to −2.21 MPa, and diminished stomatal conductance, net photosynthetic rate (from 9.13 to 2.5 μmol m^−2^ s^−1^), photosystem II photochemical efficiency (from 0.79 to 0.67), and biomass accumulation, while increased the electrolyte leakage. The shading allowed the water-deficient plants to maintain water potential (−1.58 MPa) and photosystem II efficiency (0.79) and to increase water use efficiency (from 14.80 to 86.90 μmol CO_2_/mmol H_2_O), net photosynthetic rate (from 2.40 to 9.40 μmol m^−2^ s^−1^) and biomass of leaves, crowns, and roots compared to non-shaded plants without water limitation. These results suggest that a reduction in incident light intensity attenuates the effects of stomatic and non-stomatic limitations caused by water deficit during vegetative growth of strawberry.

## Introduction

Strawberry (*Fragaria ×ananassa* Duch.) is a crop of high socio-economic value worldwide due to its organoleptic properties, contents of vitamins, minerals, and bioactive compounds such as antioxidants (Fierascu et al., 2020). Strawberry has high water requirements (Klamkowski and Treder, 2006; García-Tejero et al., 2018) and is highly susceptible to water deficit during crop establishment (El-Farhan and Pritts, 1997, Grant et al., 2012), which impacts plant growth, yield, and fruit quality (Liu et al., 2007).

In recent years, climate change has increased the frequency and intensity of drought periods worldwide (OECD 2014; Leng and Hall, 2019). Water availability is projected to decrease and global water demand to increase (OECD 2014; Mo et al. 2017), which will potentially cause unforeseen stress conditions in unprotected and even protected crops.

Plants exhibit numerous physiological, morphological, biochemical, and molecular mechanisms to cope with water deficit (Farooq et al., 2009; Ahuja et al., 2010) with variable effects on plant growth. Stomatal closure, mediated by abscisic acid, is one of the first responses to low water availability (Osakabe et al., 2014; Martin-StPaul et al., 2017) since it limits leaf transpiration; however, it also reduces CO_2_ uptake and, therefore, diminishes photoassimilate production and plant growth (Galmés et al., 2013). In strawberry plants, reduced stomatal conductance, accompanied by significantly lower photosynthetic rates, has been reported as a typical response to drought (Martínez-Ferri et al., 2016).

As water stress increases, reactive oxygen species (ROS) are produced and may cause damage to proteins, nucleic acids, lipids, and chlorophylls (Munné-Bosch and Peñuelas, 2004). ROS accumulation causes lipid peroxidation in membranes and damages in plant photosystem II (PSII; Duan et al., 2013). In strawberry, water stress has resulted in excessive generation of ROS, which increases membrane permeability and reduces PSII photochemical efficiency, *F_v_/F_m_* (Gulen et al., 2018). Consequently, reduced photosynthetic rates slow down plant growth, change the pattern of assimilate distribution (Martínez-Ferri et al., 2016), and affects the number of leaves, leaf expansion rate, total plant biomass, and yield (Blanke and Cooke, 2004; Grant et al., 2012). Depending on the cultivar, reductions between 25-37% in plant growth have been reported (Klamkowski and Treder, 2008), and decreases of 33% in yield and 17% in fruit size have been recorded (El-Farhan and Pritts, 1997).

Most herbaceous species, including strawberry, do not have efficient mechanisms to mitigate the effects of water deficit (Martínez-Ferri et al., 2016). Therefore, it is paramount to evaluate strategies aimed to improve plant performance under water-limited conditions (Ghaderi et al., 2015). The use of shading nets in environments of high light intensity and low water availability might be an effective and inexpensive technique to alleviate water deficit in strawberry (Ahemd et al., 2016). Shading has been successfully applied in many crop species including young peach (*Prunus armeniaca*) (Nicolás et al., 2005), kiwi (*Actinidia deliciosa*) (Montanaro et al., 2009), and olive (*Olea europaea*) (Sofo et al., 2009) to mitigate the effects of drought stress, and has become a common practice to improve fruit quality and plant physiology in subtropical fruits (Mditshwa et al., 2019). Likewise, in strawberry, shading nets are increasingly used, even over plastic tunnels, to protect plants and fruits from excessive sunlight (Neri et al. 2012). The positive effect of shading on plant water status and physiology is attributed to reduced intensity of incident light and changes in microclimatic variables such as air temperature, relative humidity, and vapor pressure deficit (Jifon and Syvertsen, 2003; Mditshwa et al., 2019).

In strawberry, the effects of water stress, and shading on physiological and yield responses have been independently studied (Tabatabaei et al., 2008; Casierra-Posada et al., 2012; Klamkowski et al., 2015). However, to our knowledge, there are not reports on the joint evaluation of the effects of water deficit and shading on strawberry physiology. The objective of this research was to evaluate the effect of shading on the vegetative growth of strawberry under water deficit, as a potential strategy to mitigate water shortage effects on this crop.

## Materials and Methods

### Plant Material and Growth Conditions

The study was carried out under greenhouse conditions at the Faculty of Agricultural Sciences, Universidad Nacional de Colombia, Bogotá (Colombia) located at 2640 m a.s.l. Stolons of the neutral day strawberry (*Fragaria ×ananassa*) cv. Sweet Ann (Planasa, USA) with roots and crowns of 12 and 2 cm length, respectively, were planted in 3 L plastic pots (one stolon per pot) containing a mixture of soil, blond peat, and burnt rice husk (16:1:1 w/w/w). According to the physicochemical analysis of substrate (Table S1), and nutrient requirements of strawberry (Tagliavini et al., 2005), the plants were fertilized once a week with 320 mL of a nutrient solution composed of 0.42 g L^−1^ KNO_3_, 0.86 g L^−1^ Ca(NO_3_)_2_, 0.12 g L^−1^ NH_4_H_2_PO_4_, and micronutrients (Fe, B, Mn, Zn, and Cu).

Fertilizer was distributed to avoid altering the water status of the plants according to the treatments. S and Mg were not included as part of the fertilization, due to their initial contents in the substrate (Table S1). All plants were watered maintaining the substrate water capacity (SWC) from planting until the start of the treatments. To avoid early flower development, flower buds were removed from the plants during the experiment.

Air temperature and relative air humidity were recorded with a VP-3 sensor (Decagon Devices Inc., USA) and Photosynthetically Active Radiation (PAR) was registered with PAR sensor (Vernier, USA) positioned 30 cm above the plants (Table S2); the sensors were coupled to Data Logger Decagon EM50 (Decagon Devices Inc., USA).

### Treatments

From planting, the experiment took 74 days and when plants had 3-4 expanded leaves (50 days after planting), the treatments were established in a 2 × 2 factorial design. Treatments consisted of the combination of two levels of shading and two levels of water availability: i) shading (S) with a 47% reduction in incident light (maximum and minimum diurnal PAR of 251.70 and 21.35 μmol m^−2^ s^−1^, respectively) or non-shaded (NS) conditions (maximum and minimum diurnal PAR of 476.07 and 40.28 μmol m^−2^ s^−1^, respectively) and ii) water deficit (WD) or well-watered (WW) plants. The combination of the factors resulted in four treatments: NS-WW, NS-WD, S-WW, and S-WD, and each treatment had four replicates consisting of one plant. The shading was provided with a commercially available polyethylene black net referred with a shading percentage of 33% and a weight of 31 g .m^−2^, placed 1.5 m above plants. Under shading, the average relative air humidity (RH) was 62% and the average air temperature (T) corresponded to 21 °C, while non-shaded conditions had 59% of RH and a T of 26 °C.

Based on the substrate moisture, the WW plants were maintained at a capacity of 100% Volumetric Water Content (VWC). For WD treatments, it has been reported in strawberry cultivars and wild strawberry (*Fragaria virginiana*) that water deficit during 7–15 days was enough to obtain physiological responses (O’Neill, 1983; Gulen et al., 2018). However, it has been also shown that a progressive increase in the level of stress may be valuable to detect additional changes in water relations and chlorophyll fluorescence (Liu et al., 2007; Razavi et al., 2008). Thus, WD treatments had 50% VWC during the first 15 days and, subsequently, the water supply was suspended until reaching 25% VWC, which was maintained until the end of the experiment (74 days after planting), using water content measurements as described below.

### Substrate Volumetric Water Content and Leaf Water Status

The substrate volumetric water content (SVWC) was measured daily between 9:00 and 10:00 am with a portable soil moisture sensor (ThetaProbe ML2x, Delta-T Devices, Cambridge, UK) at 10 cm depth. The SVWC is the ratio between the volume of water present and the total volume of the sample, and it is expressed as m^3^ .m^−3^. The Leaf water potential (Ψ*_lw_*) was measured in the first completely expanded leaf of the upper third of the plants at 5, 10, 17, and 24 days after treatment (DAT) at midday with a Scholander pressure chamber (PMS Instruments Model 615, CA, USA).

### Leaf Gas Exchange, Photosynthetic rate, and Water Use Efficiency

The gas exchange parameters were measured in the first completely expanded leaf of the upper third of the plants at 5, 10, 17, and 24 days after treatment (DAT). Net photosynthetic rate (*Pn*), stomatal conductance (*g_s_*), and intercellular CO_2_ concentration (*C_i_*) were registered using a photosynthesis measurement system LI-6200 (LI-COR Inc., Biosciences, USA) from 9:00 to 11:30 am with natural light (428.46 – 476.07 μmol m^−2^ s^−1^) and a CO_2_ concentration of 380 to 400 μL L^−1^. The intrinsic water use efficiency (WUEi) was calculated as the ratio of photosynthetic rate and stomatal conductance.

### Relative chlorophyll content (SPAD)

The relative chlorophyll content (SPAD values) was determined using a portable chlorophyll SPAD-502 (Konica Minolta, Sakai, Osaka, Japan) making 10 measurements on the same leaves that were used to measure photosynthetic rate.

### Chlorophyll a Fluorescence

Photosystem II photochemical efficiency (*F_v_/F_m_*) was measured in dark-adapted leaves for 30 min using a MINI-PAM modulated fluorometer (Walz^®^, GmbH Effeltrich, Germany); this measurement was performed on the same leaves used to measure photosynthetic rate. The measurements were done at a steady state of 2,000 μmol m^−2^ s^−1^ and a saturating state of 5,000 μmol m^−2^ s^−1^ of actinic light for 0.80 s.

### Membrane Permeability

Cell membrane permeability in leaves was evaluated by electrolyte leakage (EL) according to Gulen and Eris (2004). Ten 5-mm-diameter leaf discs were placed in Falcon tubes with 10 mL of deionized water at 16 °C. The electrical conductivity (EC) was determined with a conductometer (HI 9835 Hanna^®^, Spain). Electrolyte leakage values were expressed as a percentage in relation to the highest value using the equation EL = (EC_1_/EC_2_)*100, where EC_1_ = EC at 4 h, and EC_2_ = EC after heating for 15 min at 90 °C.

### Specific Leaf Area and Dry Mass Distribution

To determine the leaf area (LA) of each plant, the non-destructive estimation method was employed according to Grijalba et al. (2015), who utilized the following equation in strawberry cv. Albion: *y* = 0.8316*X*^1.9784^, where *y* is the area of the leaflet (cm^2^) and *X* is the length (cm) of each leaflet measured from its base to the apex at the central rib. From the sum of individual leaflet areas, the total leaf area of each plant was obtained. Each plant was separated into roots, leaves, and crown at the end of the experiment (24 DAT) and the plant material was dried at 65 °C until reaching a constant weight. Total dry weight and dry weights of the roots, shoots, crown, and leaves were determined, where the shoot corresponded to crown plus leaves. The specific leaf area was determined as the ratio of fresh-leaf area/dry mass.

### Statistical Analysis

The experiment consisted of a full factorial experiment with two factors (Shading and water deficit), two levels per factor, and four treatments (combinations between the levels of each factor) as detailed above. A two-way ANOVA was carried out to determine the effects of the individual factors and treatments on the analyzed variables. The data are presented for factors and their combinations. Mean comparisons were done with a Tukey’s multiple range test (*p* <0.05, 0.01, and 0.001). The statistical analysis was made using the software SAS v. 9.4 (Statistical Analysis System, USA).

## Results and Discussion

The water deficit affected all physiological variables except *C_i_*, SPAD values, SLA, RDW, and CDW, while the shading affected all variables except *C_i_* and SLA (Table 1). Significant interactions between factors (water deficit and shading) were found for Ψ*_lw_*, *g_s_*, *P_n_*, WUE_i_, *F_v_/F_m_*, EL, SLA, RDW, SDW, CDW, LDW, and TDW and are described below.

**Table 1.** Analysis of variance (ANOVA) for physiological variables of leaf water potential (Ψ*_lw_*), stomatal conductance (*g_s_*), intracellular CO_2_ concentration (*C_i_*), photosynthetic rate (*P_n_*), intrinsic water use efficiency (WUE_i_), SPAD values, photosystem II photochemical efficiency (*F_v_/F_m_*), electrolyte leakage percentage (EL), leaf area (LA), specific leaf area (SLA), shoot dry weight (SDW), root dry weight (RDW), crown dry weight (CDW), leaf dry weight (LDW), and total dry weight (TDW) of strawberry plants cv. Sweet Ann under shading and water deficit. Averages and significance are shown for each level and factor.

| Source of variation | $\Psi_{lw}$<br>(Mpa) | $g_s$<br>(mol m<sup>-2</sup> s<sup>-1</sup>) | $C_i$<br>(μmol mol<sup>-1</sup>) | $Pn$<br>(μmol m<sup>-2</sup> s<sup>-1</sup>) | $WUE_i$<br>(μmol<br>CO<sub>2</sub>/mol H<sub>2</sub>O) | SPAD<br>values | $F_v/F_m$ | EL (%) | LA<br>(cm<sup>2</sup>) | SLA<br>(cm<sup>2</sup> g<sup>-1</sup>) | RDW (g) | SDW (g) | CDW (g) | LDW (g) | TDW (g) |
| --- | --- | --- | --- | --- | --- | --- | --- | --- | --- | --- | --- | --- | --- | --- | --- |
| Well watered | -1.41 | 0.51 | 368.74 | 10.52 | 21.15 | 45.79 | 0.80 | 21.13 | 338.29 | 186.86 | 1.60 | 2.90 | 1.06 | 1.65 | 4.58 |
| Water deficit | -1.85 | 0.16 | 367.81 | 5.94 | 50.89 | 44.36 | 0.74 | 22.95 | 198.96 | 203.29 | 1.74 | 2.20 | 1.16 | 1.11 | 3.88 |
| Non-shaded | -1.87 | 0.36 | 368.06 | 5.79 | 15.56 | 42.10 | 0.74 | 22.29 | 227.60 | 191.47 | 1.37 | 2.21 | 1.01 | 1.27 | 3.68 |
| Shaded | -1.39 | 0.30 | 368.49 | 10.65 | 51.44 | 48.05 | 0.80 | 21.53 | 304.89 | 198.69 | 1.96 | 3.00 | 1.20 | 1.55 | 4.88 |
| ANOVA |  |  |  |  |  |  |  |  |  |  |  |  |  |  |  |
| Water deficit (WD) | *** | *** | n.s. | *** | *** | n.s. | *** | ** | *** | n.s. | n.s. | *** | n.s. | *** | *** |
| Shading (S) | *** | *** | n.s. | *** | *** | *** | *** | * | ** | n.s. | * | *** | * | ** | *** |
| WD x S | ** | ** | n.s. | ** | ** | n.s. | ** | ** | n.s. | ** | ** | ** | ** | * | *** |

ANOVA: \*, \*\*, and \*\*\* significantly different at the 0.05, 0.01 and 0.001 probability levels, respectively, according to the Tukey's test. n.s., not significant at $p \leq 0.05$ .

### Substrate Volumetric Water Content and Leaf Water Status

Water shortage during 24 DAT negatively affected water status in strawberry plants (Figure 1). The VWC in the WW plants varied between 0.43 and 0.40 m^3^ m^−3^. For the WD plants, the VWC was 0.22–0.21 m^3^ m^−3^ (50% SVWC) until 15 DAT and decreased to 0.13–0.10 m^3^ m^−3^ (25% SVWC) at 20 DAT, maintaining these values up to the end of the experiment (Figure 1a).

**Figure 1.**
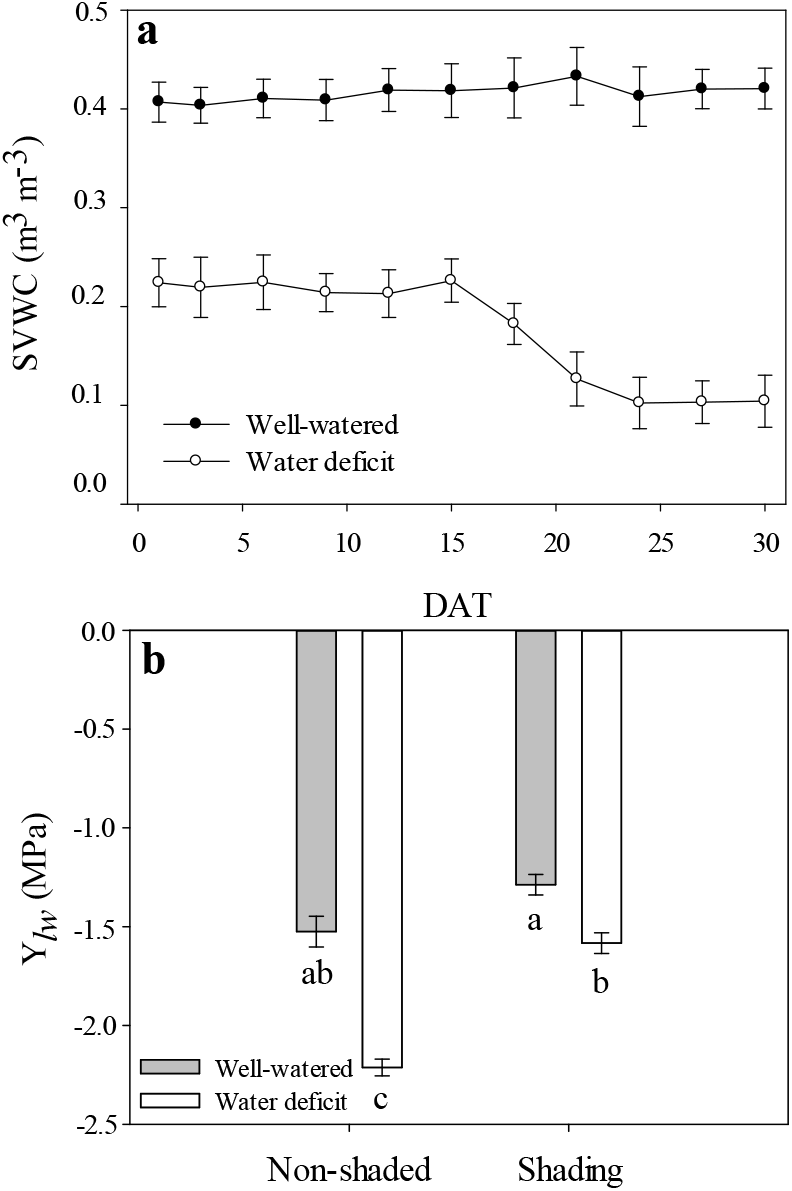
Substrate volumetric water content (SVWC) during 0-30 DAT (a) and leaf water potential (Ψ*_lw_*) at 24 DAT (b) of ‘Sweet Ann’ strawberry plants grown under non-shaded and shading conditions. DAT: days after treatment. Values are the means of four replicates, with error bars representing the standard error. Means denoted by the same letter do not significantly differ at *p* ≤ 0.01 according to the Tukey’s test.

The individual factors (water deficit and shading) and the combination of these two factors had a significant effect on leaf water potential (Ψ*_lw_*) in strawberry plants (Table 1). In the NS-WD treatment, Ψ*_lw_* was significantly lower (−2.21 MPa) than that observed in the NS-WW plants (−1.53 MPa; Figure 1b). A decrease in Ψ*_lw_* under water deficit has been reported as one of the main effects of this stress in strawberry (Klamkowski and Treder, 2008; Grant et al., 2010, 2012). The decrease in Ψ*_lw_* could be an acclimatization response to cope with adverse conditions and an indirect measure of the severity of stress (Holmgren et al., 2012). Reduced Ψ*_lw_* values might be a consequence of reduced osmotic potential, which is caused by the synthesis of compatible solutes facilitating water absorption during abiotic stress (Hasegawa et al., 2000). To counteract water deficit, strawberry plants employ osmotic adjustment, increase cell membrane stability, increase leaf thickness, and improve water use efficiency (Grant et al., 2010, 2012).

No significant differences in Ψ*_lw_* were found between the S-WW and the S-WD plants compared to the NS-WW plants (Figure 1b). These data indicate that the shading had a positive effect on the water status of water-stressed plants by avoiding reduction in Ψ*_lw_* despite the water limitation (Figure 1b). Awang and Atherton (1994) and Ma et al. (2015) reported for strawberry and other plants belonging to the Rosaceae family that shading increased Ψ*_lw_* in well-watered plants and those subjected to abiotic stress (water deficit or salinity), which, in turn, allowed a greater tolerance to the adverse effects of stress.

### Leaf Gas Exchange, Photosynthetic rate and Water Use Efficiency

The shading had a significant effect on *g_s_*, being significantly lower in the shaded plants (0.30 mol m^−2^ s^−1^) than in the non-shaded plants (0.36 mol m^−2^ s^−1^), independently of water availability (Table 1). Regarding the water deficit factor, the *g_s_* was significantly reduced from 0.509 mol m^−2^ s^−1^ in the WW plants to 0.155 mol m^−2^ s^−1^ in the WD plants. As a result of the interaction of factors (Figure 2a), the *g_s_* was significantly lower in the NS-WD, S-WW, and S-WD treatments with respect to the NS-WW (control), with the lowest values observed in the WD plants (0.17 and 0.14 mol m^−2^ s^−1^ in the NS and S treatments, respectively).

**Figure 2.**
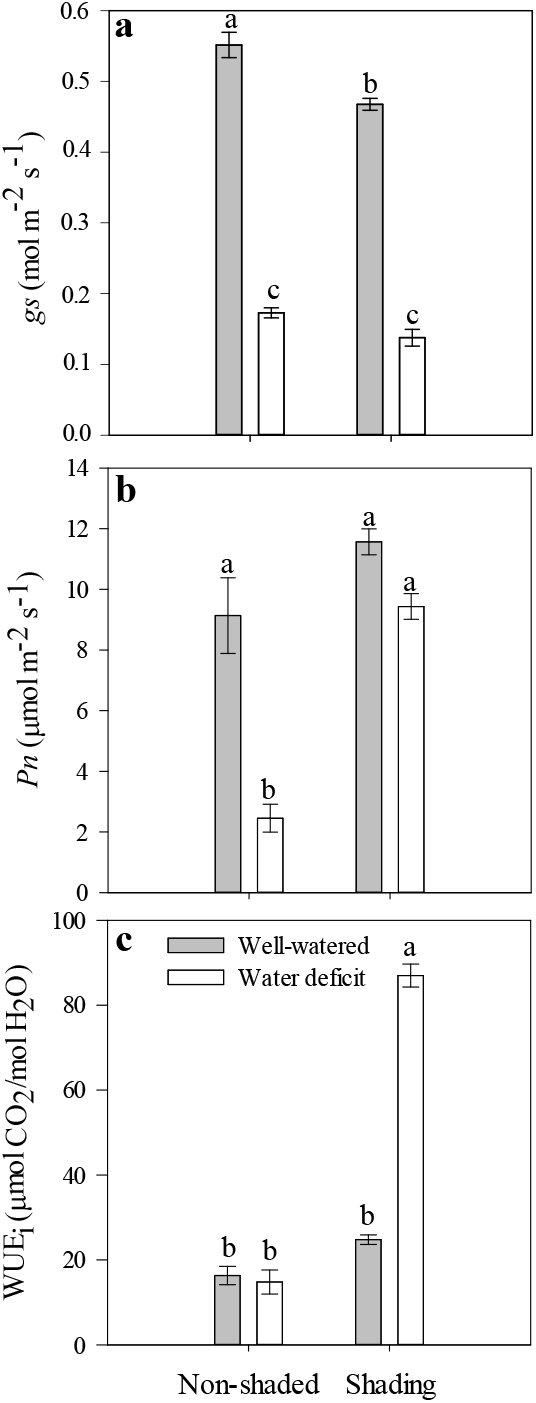
Stomatal conductance (*g_s_*) (a), net photosynthesis rate (*Pn*) (b) and intrinsic water-use efficiency (WUE_i_) (c) at 24 DAT of ‘Sweet Ann’ strawberry plants grown under non-shaded and shading conditions. DAT: days after treatment. Values are the means of four replicates, with error bars representing the standard error. Means denoted by the same letter do not significantly differ at *p* ≤ 0.01 according to the Tukey’s test.

Reductions in *g_s_* in response to shading have been reported for various species, including strawberry (Whitehead and Teskey, 1995; Gross et al., 1996; Choi et al., 2016; Gerardin et al., 2018). In shaded tobacco plants, Gerardin et al. (2018) attributed the reduced *g_s_* values to signaling mediated by the intercellular CO_2_ (*C_i_*) to optimize the concentration of CO_2_ in leaves. In this regard, Mott (1988) indicated that, in the dark, the stomata of wheat did not respond to free CO_2_ concentration and their opening depended on *C_i_*. Despite the reduction in *g_s_* caused by both water deficit and shading, the *C_i_* in strawberry leaves remained around 368 μmol mol^−1^, with no significant differences among the treatments (Table 1). Thus, the reduction in *g_s_* (Figure 2a) can be attributed to the effect of light intensity on stomata functioning. This relationship between the light intensity and stomatal density was established in wild strawberry (Jurik et al., 1982) and herbaceous plants, such as spinach (Nguyen et al., 2019). On the other hand, the S plants had a significantly higher leaf area (304.89 ± 42.33 cm^2^) than the NS plants (227.60 ± 35.81 cm^2^; Table 1), which might have affected stomatal density (Onwueme and Johnston, 2000; Qiu et al., 2018); low stomatal density in the S plants might have had a further effect on the *g_s_*.

The reduction in *g_s_* observed in the WD plants is a response that has previously been observed in many species as a strategy to prevent water loss through transpiration (Blanke and Cooke, 2004; Klamkowski and Treder, 2006). Furthermore, in strawberry plants grown under water deficit or waterlogging, a decrease in *g_s_* was observed without a reduction in *C_i_* (Blanke and Cooke, 2004; Klamkowski and Treder, 2006) as reported here. This response can be attributed to the direct effects of water stress on non-stomatal components of photosynthesis (Yordanov et al., 2000).

Shading and water availability had a significant effect on *P_n_*, being significantly lower in the WD and NS plants (Table 1). In the interaction of factors, significantly lower *P_n_* values were observed in the NS-WD plants (2.45 μmol m^−2^ s^−1^) with respect to the NS-WW plants (9.14 μmol m^−2^ s^−1^). The S-WD plants did not show significant differences in *P_n_* with the S-WW (11.57 μmol m^−2^ s^−1^) and NS-WW (9.14 μmol m^−2^ s^−1^) treatments. This indicates that shading allowed the WD plants to maintain a photosynthetic activity equal to that of the WW plants (Figure 2b) despite a significant reduction in *g_s_* in the WD-S as discussed above. The changes in microclimate might have a photoprotective effect on plants by decreasing PAR and air temperature and increasing relative humidity, thus, attenuating the negative effects of water deficit (Li et al., 2014; Ahemd et al., 2016). This could explain the significantly higher values of *P_n_* observed in the S-WD plants as compared to the NS-WD plants. In some strawberry varieties, low light intensity produced a reduction in yield and fruit quality; however, 40% shading levels did not significantly reduce plant growth or net photosynthesis (Choi et al., 2014). Our results indicated that shading at 47% did not represent a limitation for plant growth. On the contrary, this level of shading could likely reduce the effects of water deficit on *P_n_* during vegetative growth by avoiding photoinhibition.

The intrinsic water use efficiency (WUE_i_) could be indicative of plant adaptation to water-limited environments (Medrano et al., 2015). The WUE_i_ was significantly higher in the S-WD plants (86.90 μmol CO_2_/mol H_2_O) than in the other treatments (14.8 – 24.8 μmol CO_2_/mol H_2_O; Figure 2c). In strawberry plants grown under water deficit, an increase in WUE_i_ was previously found as part of plant responses to contend the water deficit (Grant et al., 2015).

The increase in WUE_i_ in the S-WD plants was because *P_n_* did not decrease, in comparison to the NS-WD plants, despite the low *g_s_* and similar *Ci* in both treatments (Table 1; Figure 2). In turn, this may be due to the non-stomatal limitations in the NS-WD plants since high solar radiation under water deficit can cause an imbalance between electron excitation and use in photosynthesis, resulting in photoinhibition (Choi et al., 2016). Moreover, a strong relationship exists between ^13^C isotope discrimination and WUE_i_ (Kumar and Singh, 2009). Grant et al. (2009) reported that strawberry plants with a low water deficit had a higher ^13^C/^12^C ratio associated with a higher WUE_i_ and increased rate of photosynthesis. In our study, a lower *g_s_*, a statistically similar *C_i_*, and the higher *P_n_* and WUE_i_ in the S-WD plants may be due to lower discrimination of ^13^C, which added to the effect of shading, might have allowed maintaining the photosynthetic activity of the S-WD plants. However, subsequent studies assessing discrimination of ^13^C under these conditions are necessary to verify this assumption.

### Relative chlorophyll content (SPAD)

The S plants had higher SPAD values (47.80 – 48.30) compared to the NS plants (41.00 – 43.30), without the influence of the water availability (Table 1; Figure 3a). The increase in SPAD values might have contributed to maintaining a high *P_n_* in the S-WD plants (Figure 2b) due to a better absorbance of incident radiation (Zhang et al., 2015). Similar results were reported in various species subjected to shading, including strawberry (Mauro et al., 2011; Luo et al., 2012; Russo and Honermeier, 2017). At the same time, Roiloa and Retuerto (2007) reported reduced levels of chlorophyll in leaves of *Fragaria vesca* grown under low light intensity. In water-deficient plants, reduced contents of chlorophyll caused by ROS have been found (Roiloa and Retuerto, 2007); however, this effect was not observed in the present experiment.

**Figure 3.**
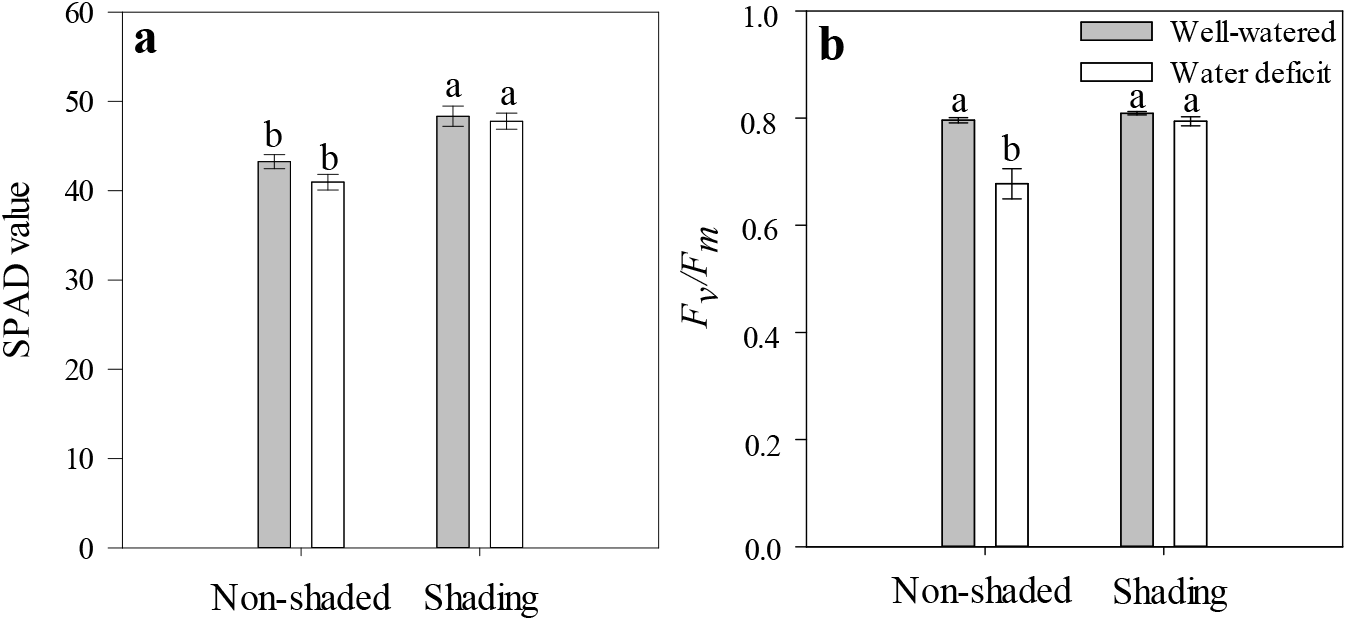
SPAD values (a) and photosystem II photochemical efficiency (*F_v_/F_m_*) (b) at 24 DAT of ‘Sweet Ann’ strawberry plants grown under non-shaded and shading conditions. DAT: days after treatment. Values are the means of four replicates, with error bars representing the standard error. Means denoted by the same letter do not significantly differ at *p* ≤ 0.01 according to the Tukey’s test.

### Chlorophyll a Fluorescence

In the NS-WD plants, *F_v_/F_m_* significantly decreased (0.68 ± 0.03) with respect to the NS-WW plants (0.80 ± 0.005), while no differences in *F_v_/F_m_* were observed between S-WD and S-WW plants (0.79 ± 0.009 vs. 0.81 ± 0.003; Figure 3b). Reduced *F_v_/F_m_* values in the NS-WD plants suggest the presence of non-stomatal limitations caused by the damage of PSII, as reported in other plants subjected to severe or moderate water deficit (Grant et al., 2010). *F_v_/F_m_* values below 0.79 are commonly reported as indicators of PSII damage by different types of stress in strawberry (Na et al., 2014; Choi et al., 2016). In this regard, in strawberry plants exposed to abiotic stress such as high temperature, *F_v_/F_m_* values from 0.72 indicated a lower PSII efficiency and were the main reason for reduced photosynthetic rates (Kadir and Sidhu, 2006). Similar results were previously found by Duan et al. (2013) in shaded strawberry plants. In this way, shading might also prevent photoinhibition in water-deficient strawberry cv. Sweet Ann (Liu et al., 2007; Zeng et al., 2010). Additionally, Choi et al. (2016) found a negative correlation between *g_s_* and *F_v_/F_m_* in strawberry leaves. In the present study, reduced *g_s_* values in shaded plants might be related to the PSII efficiency, which allowed the S-WD plants to tolerate stress conditions and maintain their photosynthetic activity.

### Membrane Permeability

The EL was significantly affected by shading, water limitation, and the combination of both factors (Table 1). In the NS-WD plants, the EL (24.5%) was significantly higher compared to the NS-WW (20.6%), while no differences were observed in EL between the NS-WW, S-WD, and S-WW plants (Figure 4). Higher EL values in the NS-WD treatment might indicate cell membrane damage due to a high activity of ROS that caused lipid peroxidation and loss of selective permeability of membranes (McDonald and Archbold, 1998; Sun et al., 2015). These results suggest that a lower light intensity favored the membrane integrity by decreasing ROS production, attenuating the effects of water stress on strawberry plants.

**Figure 4.**
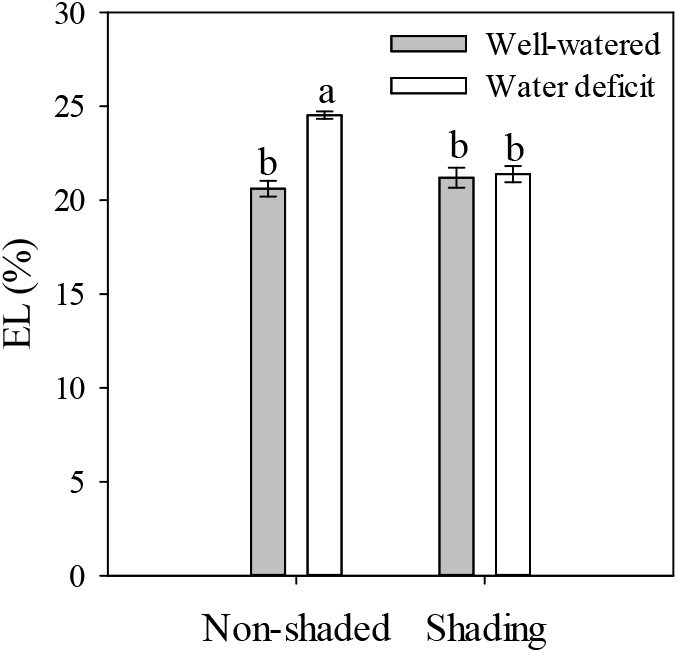
Electrolyte leakage (EL) at 24 DAT of ‘Sweet Ann’ strawberry plants grown under non-shaded and shading conditions. DAT: days after treatment. Values are the means of four replicates, with error bars representing the standard error. Means denoted by the same letter do not significantly differ at *p* ≤ 0.01 according to the Tukey’s test.

### Specific Leaf Area and Dry Mass Distribution

The water deficit and shading had a significant effect on the accumulation of dry mass in strawberry plants. For the specific leaf area (SLA), a significant increase was observed only in the NS-WD treatment (218.5 cm^2^ g^−1^) compared to the NS-WW plants (171.2 cm^2^ g^−1^), while the other treatments did not differ in SLA (Figure 5a). The root dry weight (RDW) was higher in the S-WD plants (2.3 g) with respect to the NS-WW and NS-WD plants (1.2–1.5 g), but without differences with the S-WW plants (Figure 5b). The shoot dry weight (SDW) was significantly lower in the NS-WD plants (1.70 g) in comparison to the other treatments (2.70 – 3.10 g) that did not differ in this variable (Figure 5c). The S-WD plants accumulated the highest crown weight, CDW (1.40 g) without significant differences with the NS-WW plants, while the S-WW and NS-WD plants had the lowest values of CDW (0.90–1.0 g) without differences with the NS-WW (1.1 g) (Figure 5d). The leaf dry weight (LDW) was significantly lower in the NS-WD (0.8 g) as compared to the other treatments (1.40–1.70 g), which did not differ in this variable (Figure 5e). The highest total dry weights (TDW) were registered in the S-WW (4.80 g) and S-WD (4.90 g) compared to the NS-WW (4.20 g); on the contrary, the NS-WD plants had the lowest TDW (2.90 g; Figure 5f).

**Figure 5.**
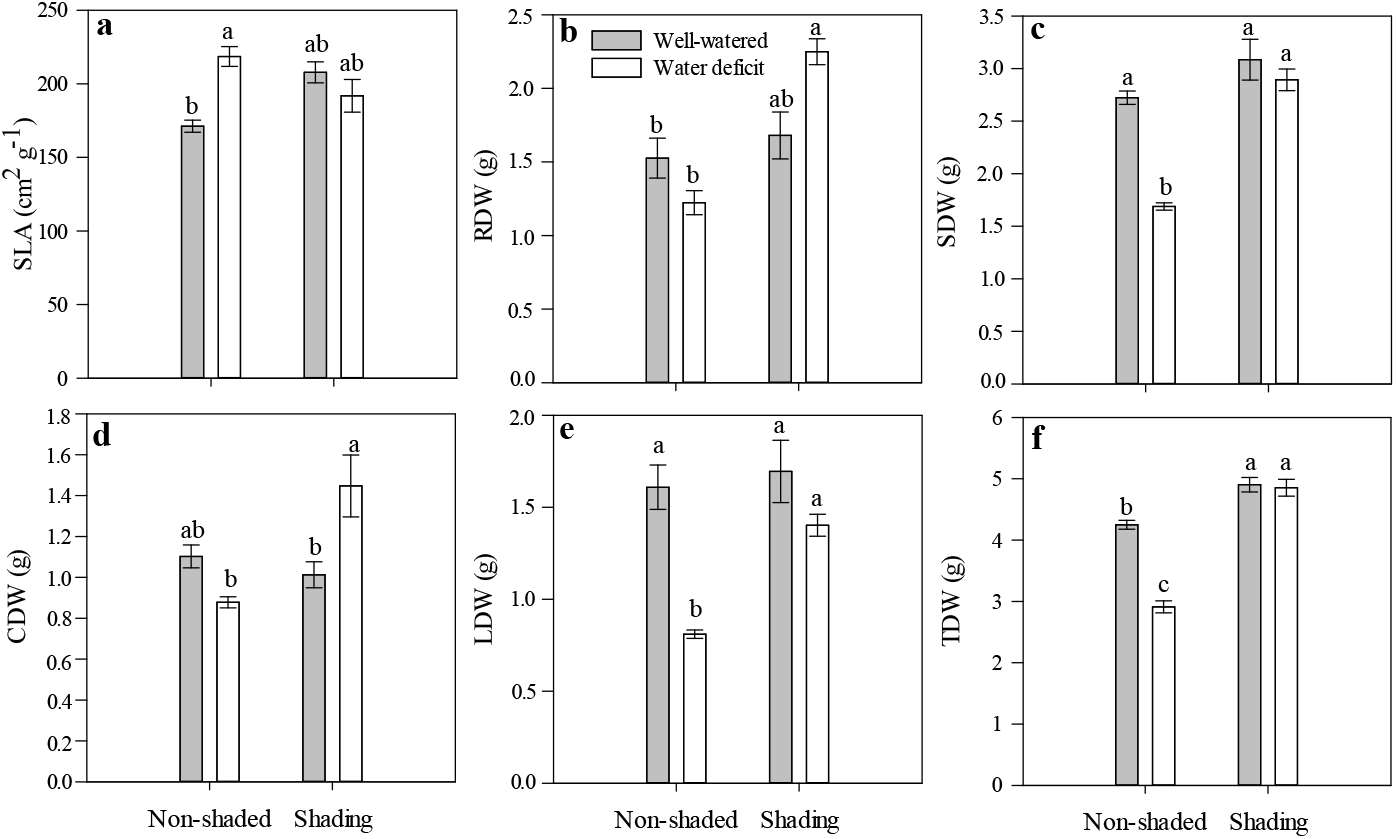
Specific leaf area (SLA) (a), root dry weight (RDW) (b), shoot dry weight (SDW) (c), crown dry weight (CDW) (d), leaves dry weight (LDW) (e), and total dry weight (TDW) (f) at 24 DAT of ‘Sweet Ann’ strawberry plants grown under non-shaded and shading conditions. DAT: days after treatment. Values are the means of four replicates, with error bars representing the standard error. Means denoted by the same letter do not significantly differ at P ≤ 0.01 according to the Tukey’s test. For LDW differences are given at *p* ≤ 0.05.

An increase in SLA in the NS-WD plants was apparently related to a reduced leaf thickness and can be attributed to the reduction in leaf area and an imbalance between assimilation, distribution, and use of carbon during water deficit (Álvarez et al., 2011). In fact, the WD plants had significantly less leaf area (198.90 ± 15.11 cm^2^, *p* < 0.001) compared to the WW plants (338.29 ± 29.76 cm^2^; Table 1) and a reduced *P_n_* in the NS-WD treatment (Figure 2b). Leaf area reduction is a typical response to water deficit to reduce transpiration area and is accompanied by reduced carbon assimilation due to stomatal and non-stomatal limitations. These responses restrict plant growth and were observed in the lower weight of the NS-WD plants (Figure 5).

Another factor that could influence the accumulation of biomass under water stress is the increase in the use of carbohydrates for maintenance respiration of existing organs, due to the drastic decrease in photosynthetic rate (Cameron et al., 1999; Sánchez-Blanco et al., 2009). However, further studies would be necessary to assess the respiration of strawberry plants under water deficit and shading.

In the S-WD plants, shading favored a better carbon balance by maintaining and increasing dry weight in all organs compared to the NS-WD plants (Figure 5b-d). A higher accumulation of biomass was related to a greater *Pn*, WUE_i_, and *F_v_/F_m_* in the S-WD, compared with the NS-WD plants, which indicates attenuation of the effects of water deficit in shaded plants and agrees with previous reports on the effects of low radiation under stress conditions (Montanaro et al., 2009). The highest accumulation of photoassimilates in roots and crowns of the S-WD plants (Figure 5b-d) may represent an advantage for plant establishment under water deficit or during recovery after a period of water deficit. In strawberry, favorable effects on the development of vegetative and reproductive buds have been reported depending on the amount and type of carbohydrates accumulated in crowns (Kirschbaum et al., 2010).

## Conclusions

The reduction of incident light by 47% generated a microclimate that mitigated the effect of stomatic and non-stomatic limitations in strawberry plants cv. Sweet Ann under water deficit. By reducing photosynthetically active radiation, shading induced a better water balance and a higher water use efficiency. Shading improved photosynthetic performance and increased biomass accumulation in water-deficient plants. In the present study, the use of shading nets has proven to be an effective alternative to manage water stress during vegetative growth. Future research assessing its potential and effects during strawberry production would be valuable to incorporate it as a common practice in this crop.

## Author’s contribution

H.A. Cordoba-Novoa, M.M. Pérez-Trujillo, and B.E. Cruz designed and performed the experiments and analyzed the data. L.P. Moreno, S. Magnitskiy, and N. Flórez guided the research and provided technical support for designing and conducting the experiments. H.A. Cordoba-Novoa, S. Magnitskiy, and L.P. Moreno wrote the manuscript. All authors read and approved the final manuscript.

